## Supplementary material for "Automated analysis of small RNA datasets with RAPID": Files to reproduce the analysis done in manuscript: Fig2.R

```
library("reshape2")
library("ggplot2")
library("viridis")
library("scales")
library("knitr")
library("gplots")
library("RColorBrewer")

COSMO = theme_bw() + theme(plot.title = element_text(size = 16, hjust = 0.5), 
                           axis.title.x = element_text(size = 16), 
                           axis.title.y = element_text(size = 16), 
                           axis.text.x = element_text(size = 14), 
                           axis.text.y = element_text(size = 14), 
                           legend.title = element_text(size = 14), 
                           legend.text = element_text(size = 14),
                           strip.text = element_text(face="bold", size=14))


allNames=c("ETS", "ITS1", "ITS2", "r17S", "r28S", "r58S")

tt=read.table("./rDNA_51A/rDNAClusterRegions/alignedReads.sub.compact",stringsAsFactors=FALSE)
tt=tt[which(tt$V1 %in% allNames),]
tt_a=melt(table(tt$V4,tt$V2, tt$V1))
tt_a$sero="51A"

tt=read.table("./rDNA_51B/rDNAClusterRegions/alignedReads.sub.compact",stringsAsFactors=FALSE)
tt=tt[which(tt$V1 %in% allNames),]
tt_b=melt(table(tt$V4,tt$V2, tt$V1))
tt_b$sero="51B"

tt=read.table("./rDNA_51D/rDNAClusterRegions/alignedReads.sub.compact",stringsAsFactors=FALSE)
tt=tt[which(tt$V1 %in% allNames),]
tt_d=melt(table(tt$V4,tt$V2, tt$V1))
tt_d$sero="51D"

tt=read.table("./rDNA_51H/rDNAClusterRegions/alignedReads.sub.compact",stringsAsFactors=FALSE)
tt=tt[which(tt$V1 %in% allNames),]
tt_h=melt(table(tt$V4,tt$V2, tt$V1))
tt_h$sero="51H"

tt_all=rbind(tt_a,tt_b,tt_d,tt_h)
tt_all$Var3=gsub("r17S","17S",tt_all$Var3)
tt_all$Var3=gsub("r58S","5.8S",tt_all$Var3)
tt_all$Var3=gsub("r28S","25S",tt_all$Var3)
tt_all$Var3_order=factor(tt_all$Var3, levels = c("ETS","17S","ITS1","5.8S","ITS2","25S"))
p=ggplot(tt_all, aes(x=Var2, y=value, fill=Var1))+
    geom_bar(stat = "identity")+xlab("Read length")+ylab("Read count (log10)") +
    facet_grid(sero~Var3_order,scales = "free")+
    scale_y_continuous(trans = log10_trans())+
    guides(fill=guide_legend(title="Strand")) +
    scale_fill_viridis(discrete = T, option = "D") +
    COSMO
ggsave("./Fig2.pdf",p, width = 11.69, height = 8.27, units = "in")
```
