## Supplementary material for "Automated analysis of small RNA datasets with RAPID": Files to reproduce the analysis done in manuscript: Fig3.R

```
library("reshape2")
library("ggplot2")
library("scales")
library("knitr")
library("gplots")
library("RColorBrewer")
library("viridis")

####FUNCTIONS####
calcCOV <- function(df){
  df2=df
  for(i in 1:nrow(df)){ df2$COV[i]=round(sd(as.numeric(df[i,-1]))/(mean(as.numeric(df[i,-1]))),digits = 2)}
  return(df2)
}

calcRLE <-function(df){
  df2=df
  df2$COV=NULL
  for(i in 1:nrow(df2)){
    medVal=median(as.numeric(df[i,2:7]))
    for(j in 2:ncol(df2)){
      df2[i,j]=round(log((1+df[i,j])/(1+medVal)),digits = 2)
      #df2[i,j]=round(log(df[i,j]/medVal),digits = 2)
    }
  }
  return(df2)
}

####DONE FUNCTIONS####

COSMO = theme_bw() + theme(plot.title = element_text(size = 16, hjust = 0.5), 
                           axis.title.x = element_text(size = 18), axis.title.y = element_text(size = 16), 
                           axis.text.x = element_text(size = 18), axis.text.y = element_text(size = 16), 
                           legend.title = element_text(size = 18), legend.text = element_text(size = 16),
                           panel.border = element_blank(), 
                           panel.grid.major = element_blank(),
                           panel.grid.minor = element_blank(),
                           axis.line = element_line(colour = "black"))

ex2bgc=read.table("./exbgc_NormalizedValues.dat", header = T, stringsAsFactors = F)
ex2wobgc=read.table("./exwobgc_NormalizedValues.dat", header = T, stringsAsFactors = F)
ex2DES=read.table("./exdes_NormalizedValues.dat", header = T, stringsAsFactors = F)

####Coeff of Variation - Example genes
ex2bgc_rdCnt = calcCOV(as.data.frame(dcast(ex2bgc, region~samples, value.var = 'readCounts')))
ex2bgc_rdCnt_Norm = calcCOV(as.data.frame(dcast(ex2bgc, region~samples, value.var = 'readCountsNorm')))
ex2_rdCnt_wobgc_Norm = calcCOV(as.data.frame(dcast(ex2wobgc, region~samples, value.var = 'readCountsNorm')))
ex2_rdCnt_NormDES = calcCOV(as.data.frame(dcast(ex2DES, region~samples, value.var = 'readCountsNorm')))

ex2COVs=as.data.frame(cbind(ex2bgc_rdCnt$region, ex2bgc_rdCnt$COV, ex2bgc_rdCnt_Norm$COV, ex2_rdCnt_wobgc_Norm$COV, ex2_rdCnt_NormDES$COV))
colnames(ex2COVs)=c("Region","Raw", "KDCS", "TCS", "DESeq2")
ex2COVs$Region=c("1","2","3","4")
ex2COVs_melted=melt(ex2COVs,id.vars = "Region",variable.name = "Method", value.name = "CoV")
ex2COVs_melted$CoV=as.numeric(as.character(ex2COVs_melted$CoV))

pdf("./Fig3.pdf")
print(ggplot(ex2COVs_melted, aes(x = Region, y = CoV, color=Method, group=Method))+
        geom_point()+geom_line()+
        theme_bw()+
        theme(axis.text.x = element_text(angle = 90, hjust = 1, vjust = 0.45))+
        labs(x="Example sRNA region", y="Coefficient of variation \n (read counts)",title="")+
        guides(color=guide_legend(title="Normalization \nmethod"))+
        scale_color_viridis(discrete = T, option = "D")+
        COSMO
)
dev.off()
```
