## Supplementary material for "Automated analysis of small RNA datasets with RAPID": Files to reproduce the analysis done in manuscript: Fig4.R

```
library("reshape2")
library("scales")
library("gplots")
library("RColorBrewer")

args <- commandArgs(trailingOnly = TRUE)
df=read.table("./spombe_Allnorm/NormalizedValues.dat", header = T, stringsAsFactors = F)
df$ASratioNormLog = log2(1+df$ASratioNorm)

rdCntNormLog=as.data.frame(dcast(df, region~samples, value.var = 'ASratioNormLog'))
rownames(rdCntNormLog)=rdCntNormLog$region
rdCntNormLog$region=NULL

#cpalette<-colorRampPalette(c("red", "yellow", "green"))(n = 92)
cpalette<-colorRampPalette(c("white","green","green4","violet","purple"))(n=92)
pdf("./Fig4.pdf")
heatmap.2(as.matrix(rdCntNormLog), col=cpalette, density.info = "none", trace = "none", keysize = 1, margin =c(5,5), 
          key.xlab = "AS ratio", dendrogram = "both", cexCol = 1.2, labRow = "" )
dev.off()
```
