## Supplementary material for "Automated analysis of small RNA datasets with RAPID": Files to reproduce the analysis done in manuscript: ReproducePlots.sh

```
##################################################################################################
#												 #
# Supplementary script for the manuscript "Automated analysis of small RNA datasets with RAPID"  #
#				Karunanithi et. al.,						 #
#												 #
#   This script assumes you have the following softwares installed and in PATH:			 #
#	conda											 #
#	cutadapt (v1.8.1)									 #
#	trim_galore										 #
#	RAPID as a condo env (see https://rapid-doc.readthedocs.io/en/latest/Installation.html)	 #
#												 #
#   This script reproduces the analysis done in this manuscript and it 				 #
#   reproduces figures (except Fig.1). 								 #
#												 #
#												 #
#   RUN THE SCRIPT IN THE FOLDER WHERE YOU HAVE ALL ASSOCIATED SUPPLEMENTARY FILES DOWNLOADED	 #
#												 #
#   UPDATE THE CONDA ENVIRONMENT VARIABLE FOR RAPID BELOW					 #
#												 #
##################################################################################################
 
ENV_NAME=<environment_name>

####################################################### Data Download, and Preprocessing

### Paramecium tetraurelia related
wget --no-passive-ftp ftp.sra.ebi.ac.uk/vol1/fastq/ERR250/000/ERR2503570/ERR2503570.fastq.gz
wget --no-passive-ftp ftp.sra.ebi.ac.uk/vol1/fastq/ERR250/001/ERR2503571/ERR2503571.fastq.gz
wget --no-passive-ftp ftp.sra.ebi.ac.uk/vol1/fastq/ERR250/002/ERR2503572/ERR2503572.fastq.gz
wget --no-passive-ftp ftp.sra.ebi.ac.uk/vol1/fastq/ERR250/003/ERR2503573/ERR2503573.fastq.gz
wget --no-passive-ftp ftp.sra.ebi.ac.uk/vol1/fastq/ERR250/004/ERR2503574/ERR2503574.fastq.gz
wget --no-passive-ftp ftp.sra.ebi.ac.uk/vol1/fastq/ERR250/005/ERR2503575/ERR2503575.fastq.gz
wget --no-passive-ftp ftp.sra.ebi.ac.uk/vol1/fastq/ERR250/006/ERR2503576/ERR2503576.fastq.gz
wget --no-passive-ftp ftp.sra.ebi.ac.uk/vol1/fastq/ERR250/007/ERR2503577/ERR2503577.fastq.gz


# Merge replicates, and rename samples
cat ERR2503570.fastq.gz ERR2503571.fastq.gz >51A_unfilt.fq.gz
cat ERR2503572.fastq.gz ERR2503573.fastq.gz >51B_unfilt.fq.gz
cat ERR2503574.fastq.gz ERR2503575.fastq.gz >51D_unfilt.fq.gz
cat ERR2503576.fastq.gz ERR2503577.fastq.gz >51H_unfilt.fq.gz

# Data trimming
# The reads are already adapter trimmed.

# Filter Reads (you need install cutadapt(v1.8.1) to perform this)
cutadapt -m 21 -M 25 ./51A_unfilt.fq.gz -o ./51A.fq.gz
cutadapt -m 21 -M 25 ./51B_unfilt.fq.gz -o ./51B.fq.gz
cutadapt -m 21 -M 25 ./51D_unfilt.fq.gz -o ./51D.fq.gz
cutadapt -m 21 -M 25 ./51H_unfilt.fq.gz -o ./51H.fq.gz

#Remove raw and intermediate files
rm *_unfilt.fq.gz ERR250357*.fastq.gz


### Spombe related
wget --no-passive-ftp ftp.sra.ebi.ac.uk/vol1/fastq/SRR444/006/SRR4449666/SRR4449666.fastq.gz 
wget --no-passive-ftp ftp.sra.ebi.ac.uk/vol1/fastq/SRR444/007/SRR4449667/SRR4449667.fastq.gz 
wget --no-passive-ftp ftp.sra.ebi.ac.uk/vol1/fastq/SRR444/008/SRR4449668/SRR4449668.fastq.gz 
wget --no-passive-ftp ftp.sra.ebi.ac.uk/vol1/fastq/SRR444/009/SRR4449669/SRR4449669.fastq.gz
cat SRR4449* >WT_24h.fq.gz

wget --no-passive-ftp ftp.sra.ebi.ac.uk/vol1/fastq/SRR444/000/SRR4449690/SRR4449690.fastq.gz 
wget --no-passive-ftp ftp.sra.ebi.ac.uk/vol1/fastq/SRR444/001/SRR4449691/SRR4449691.fastq.gz 
wget --no-passive-ftp ftp.sra.ebi.ac.uk/vol1/fastq/SRR444/002/SRR4449692/SRR4449692.fastq.gz 
wget --no-passive-ftp ftp.sra.ebi.ac.uk/vol1/fastq/SRR444/003/SRR4449693/SRR4449693.fastq.gz 
cat SRR4449* >ago1D_24h.fq.gz

wget --no-passive-ftp ftp.sra.ebi.ac.uk/vol1/fastq/SRR444/004/SRR4449714/SRR4449714.fastq.gz 
wget --no-passive-ftp ftp.sra.ebi.ac.uk/vol1/fastq/SRR444/005/SRR4449715/SRR4449715.fastq.gz 
wget --no-passive-ftp ftp.sra.ebi.ac.uk/vol1/fastq/SRR444/006/SRR4449716/SRR4449716.fastq.gz 
wget --no-passive-ftp ftp.sra.ebi.ac.uk/vol1/fastq/SRR444/007/SRR4449717/SRR4449717.fastq.gz 
cat SRR4449* >clr4D_24h.fq.gz

wget --no-passive-ftp ftp.sra.ebi.ac.uk/vol1/fastq/SRR444/008/SRR4449738/SRR4449738.fastq.gz 
wget --no-passive-ftp ftp.sra.ebi.ac.uk/vol1/fastq/SRR444/009/SRR4449739/SRR4449739.fastq.gz 
wget --no-passive-ftp ftp.sra.ebi.ac.uk/vol1/fastq/SRR444/000/SRR4449740/SRR4449740.fastq.gz 
wget --no-passive-ftp ftp.sra.ebi.ac.uk/vol1/fastq/SRR444/001/SRR4449741/SRR4449741.fastq.gz
cat SRR4449* >dcr1D_24h.fq.gz 

# Adapter trimming, and Read filtering (you need install cutadapt(v1.8.1) to perform this)
for file in *.fq.gz; 
do 
name=`echo $file|cut -d. -f1`; 
echo ${name}_trimmed.fq.gz;
trim_galore --length 18 --max_length 25 $file
mv ${name}_trimmed.fq.gz $file
done;

#Remove raw and intermediate files
rm SRR4449*.fastq.gz *_trimming_report.txt

### Move in to conda environment
source activate $ENV_NAME

# Build index for rDNA (bowtie2 installation necessary)
bowtie2-build rDNAcluster.fa rDNAIndex
bowtie2-build Spombe_ASM294v2.fasta Spombe_ASM294v2


#Run the basic statistics for Paramecium data.
rapidStats.sh -o=./rDNA_51A/ -f=./51A.fq.gz -a=./rDNAClusterRegions.bed -i=./rDNAIndex
rapidStats.sh -o=./rDNA_51B/ -f=./51B.fq.gz -a=./rDNAClusterRegions.bed -i=./rDNAIndex
rapidStats.sh -o=./rDNA_51D/ -f=./51D.fq.gz -a=./rDNAClusterRegions.bed -i=./rDNAIndex
rapidStats.sh -o=./rDNA_51H/ -f=./51H.fq.gz -a=./rDNAClusterRegions.bed -i=./rDNAIndex

#Run the normalisation to compare all serotypes.
rapidNorm.sh -o=./rDNA_Allnorm/ -c=./rDNA_allsamples.config -a=./rDNAClusterRegions.bed

#Run the basic statistics for every Pombe data.
rapidStats.sh -o=./WT_24h/ -f=./WT_24h.fq.gz -a=./sRNAenrichedGenelist_mmc2.bed -i=./Spombe_ASM294v2
rapidStats.sh -o=./ago1D_24h/ -f=./ago1D_24h.fq.gz -a=./sRNAenrichedGenelist_mmc2.bed -i=./Spombe_ASM294v2
rapidStats.sh -o=./clr4D_24h/ -f=./clr4D_24h.fq.gz -a=./sRNAenrichedGenelist_mmc2.bed -i=./Spombe_ASM294v2
rapidStats.sh -o=./dcr1D_24h/ -f=./dcr1D_24h.fq.gz -a=./sRNAenrichedGenelist_mmc2.bed -i=./Spombe_ASM294v2

#Run the normalisation to compare all serotypes.
rapidNorm.sh -o=./spombe_Allnorm/ -c=./all_only24h_sRNA.config -a=./sRNAenrichedGenelist_mmc2.bed

######## Reproduce Figures 2, 3, and 4 in the Manuscript ##########

Rscript Fig2.R
Rscript Fig3.R
Rscript Fig4.R

######## All Visualisation reports for the data analysed (One sample output is created for each case study; uncomment others and run only if necessary, as it might take considerable time) #########

#To create all the comparison plots for Paramecium

rapidVis.sh -t=compare -o=./rDNA_Allnorm/

#To create all the plots - Basic statistics of each Paramecium sample

rapidVis.sh -t=stats -o=./rDNA_51A/rDNAClusterRegions/ -a=./rDNAClusterRegions.bed 
#rapidVis.sh -t=stats -o=./rDNA_51B/rDNAClusterRegions/ -a=./rDNAClusterRegions.bed 
#rapidVis.sh -t=stats -o=./rDNA_51D/rDNAClusterRegions/ -a=./rDNAClusterRegions.bed 
#rapidVis.sh -t=stats -o=./rDNA_51H/rDNAClusterRegions/ -a=./rDNAClusterRegions.bed 

#To create all the comparison plots for Pombe
rapidVis.sh -t=compare -o=./spombe_Allnorm/

#To create all the plots - Basic statistics of each Pombe sample
rapidVis.sh -t=stats -o=./WT_24h/sRNAenrichedGenelist_mmc2/ -a=./sRNAenrichedGenelist_mmc2.bed 
#rapidVis.sh -t=stats -o=./ago1D_24h/sRNAenrichedGenelist_mmc2/ -a=./sRNAenrichedGenelist_mmc2.bed 
#rapidVis.sh -t=stats -o=./clr4D_24h/sRNAenrichedGenelist_mmc2/ -a=./sRNAenrichedGenelist_mmc2.bed 
#rapidVis.sh -t=stats -o=./dcr1D_24h/sRNAenrichedGenelist_mmc2/ -a=./sRNAenrichedGenelist_mmc2.bed 


source deactivate
```
